# Pancreatlas™: applying an adaptable framework to map the human pancreas in health and disease

**DOI:** 10.1101/2020.03.27.006320

**Authors:** Diane C. Saunders, James Messmer, Irina Kusmartseva, Maria L. Beery, Mingder Yang, Mark A. Atkinson, Alvin C. Powers, Jean-Philippe Cartailler, Marcela Brissova

## Abstract

Human tissue phenotyping generates complex spatial information from numerous imaging modalities, yet images typically become static figures for publication and original data and metadata are rarely available. While comprehensive image maps exist for some organs, most resources have limited support for multiplexed imaging or have non-intuitive user interfaces. Therefore, we built a Pancreatlas™ resource that integrates several technologies into a novel interface, allowing users to access richly annotated web pages, drill down to individual images, and deeply explore data online. The current version of Pancreatlas contains over 800 unique images acquired by whole-slide scanning, confocal microscopy, and imaging mass cytometry, and is available at https://www.pancreatlas.org. To create this human pancreas-specific biological imaging resource, we developed a React-based web application and Python-based application programming interface, collectively called Flexible Framework for Integrating and Navigating Data (FFIND), which can be adapted beyond Pancreatlas to meet countless imaging or other structured data management needs.

## Introduction

Rapid advances in microscopy, live cell imaging, and multiplexing technologies are generating a wealth of rich and increasingly complex imaging data, creating an enormous challenge to organize, process, and share this data in a way that facilitates meaningful scientific advancements. Scientists traditionally observe image data as static figures in publications or, when available online, in formats limited to only three color channels with little to no interactivity. Furthermore, comprehensive datasets encompass images generated by multiple imaging platforms and modalities, which are often acquired in proprietary formats and require individualized, proprietary image browsers that are difficult to integrate into a web environment. Therefore, there is an unmet need for more sophisticated image management and dissemination systems with capabilities to integrate data across different imaging platforms.

Although the demand for better image management solutions is apparent, software development is inherently difficult and expensive. Moreover, those who stand to benefit most from imaging data and their interpretation are usually not software developers, but rather scientists investigating complex biology and disease. While the clinical imaging community has benefited from software and database solutions driven by advances in patient care, the basic science community still relies heavily on non-enterprise level software that is developed in-house and chronically under-funded. This poses significant challenges for the basic science community to share and access imaging data in a way that is “biologist-friendly,” scalable, and that leverages existing technology.

Our research team focused on this challenge from the perspective of creating an online resource to document pancreatic architecture over the human lifespan. No reference datasets are available for human pancreas development, unlike other organ systems^1–5^, and this knowledge gap is quite limiting for those working toward an understanding of diabetes, pancreatitis, and pancreatic cancer. We assembled a multidisciplinary team of bioinformatics specialists, software developers, and biologists to build an “atlas” of the human pancreas. We chose to integrate existing tools and workflows wherever possible, layering multiple systems to meet project-specific needs and systematically cataloging the process. We prioritized the ability to handle images with more than 30 channels, allowing display of individual cell markers in user-specified combinations while still preserving spatial relationships within the context of the entire tissue section. During the process, we realized that our modular approach could not only be repurposed to share image collections in other fields of research, but it could also be adapted to organize virtually any data content or type in a streamlined website. Thus, Pancreatlas represents just one implementation of our generalized framework for displaying datasets, Flexible Framework for Integrating and Navigating Data (FFIND).

The FFIND platform and its Pancreatlas implementation are centered on the principles of automation, scalability, configuration, and simplicity for end-users. By housing data in well-defined collections, we provide curated points of entry to vast amounts of data; our filtering menus allow users to view and refine data based on multiple variables of interest. Additionally, we have integrated metadata annotations to offer information about human samples and to encourage adoption of standardized, field-specific nomenclature. By connecting images from several independent programs and facilitating exploration of imaging data from high-impact publications, Pancreatlas has the potential to accelerate understanding of human pancreas biology, integrate data from other fields such as cystic fibrosis and pancreatic cancer, and lead to transformative changes in diabetes care.

## Results

### Flexible Framework for Integrating and Navigating Data (FFIND)

Underlying Pancreatlas is FFIND, a generalized framework we developed to organize data and metadata originating from diverse sources, and then publish it via a user interface that prioritizes data exploration and discovery. FFIND enhances the value of existing databases, filesystems, and/or software by streamlining data retrieval and delivery. It is composed of both (1) a server-side (back-end) application programming interface (API), which links and retrieves data from different sources, and (2) a client-side (front-end) web application, which provides a data-agnostic, modern user interface (**Figure 1A**).

**Figure 1.**
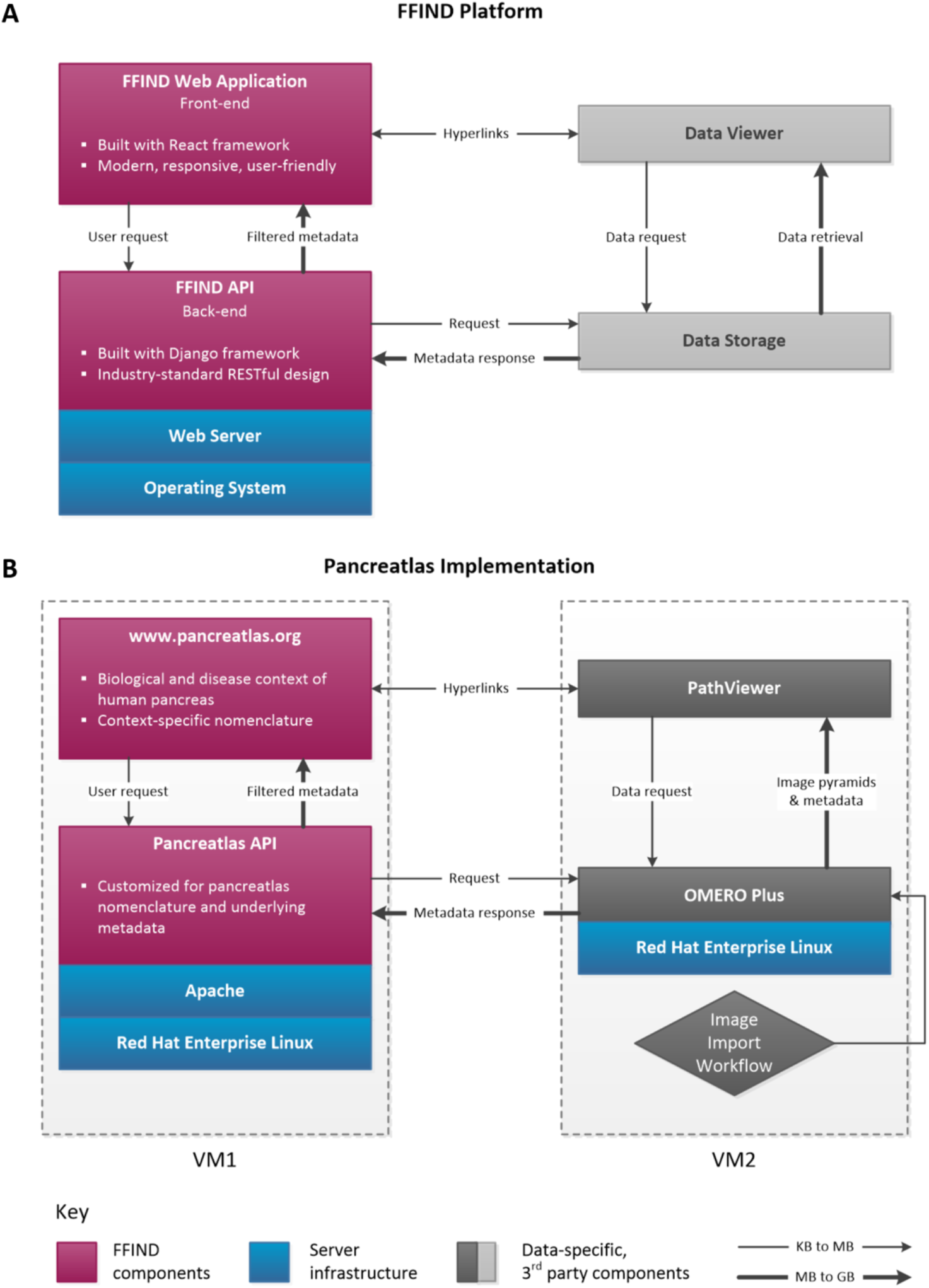
FFIND platform and its Pancreatlas implementation. Schematic showing general connectivity and data flow across (A) FFIND platform and (B) Pancreatlas implementation. In both panels, novel components are depicted in magenta, shown in relationship to server infrastructure (blue) and 3rd party components (grey). The thin arrows represent small data requests and large arrows represent larger data responses. **(A)** The intuitive web application of FFIND allows end users (researchers, clinicians, biologists) to seamlessly browse modular datasets annotated with field-specific metadata, and it can also launch existing data visualization clients to provide a cohesive browsing experience. The FFIND application programming interface (API) connects the web application to an underlying data server or storage component, which can be configured according to project needs. **(B)** To create Pancreatlas, the OMERO Plus server was installed on a virtual machine (VM2) for image management. Data and associated metadata were loaded directly from file stores and proprietary imaging servers (not pictured here), and the web application was built to retrieve and display images in PathViewer, the web client associated with the OMERO Plus server.

The FFIND back-end API is built atop of the Python-based Django Web and Django Representational State Transfer (REST) frameworks, which allowed for rapid application development using well-established design patterns and idioms, including implementing a RESTful API and caching of frequently accessed data in order to improve performance and user experience.

FFIND’s front-end web application was developed with the Javascript-based React framework. Associated dependencies include Reactstrap (port of the popular Bootstrap framework to React), tinycolor2 (used to manipulate background colors), and axios (used to communicate with APIs). Numerous components were developed that specify user interface elements, business logic, data retrieval, and interactivity. Key components of the application include the grid (default) view for displaying a table of data objects (*e.g*., images and associated descriptors in the case of Pancreatlas), preview cards for displaying a focused view of a single data object and its descriptors, and a filter panel that allows users to eliminate non-relevant objects from view in real time. In addition, a matrix view provides an intuitive mechanism for specifying the intersect of two metadata attributes and returning matching data objects in the context of the grid view. Furthermore, a time-series component was developed for age group selection, based on annotated age data per object.

### Pancreatlas as an FFIND Implementation

Pancreatlas is a domain- and organ-specific implementation of FFIND (**Figure 1B**) where data objects are images. The implementation includes functionality to retrieve and package data from Open Microscopy Environment Remote Objects Plus (OMERO Plus, Glencoe Software) using the omero-py library. OMERO Plus provides controlled access to imaging data and metadata through tiered databases, middleware, and remote client applications, allowing it to function as both a project data management tool and image data publication system^6,7^. FFIND’s API connects to the OMERO Plus API to retrieve data and metadata for individual images and image collections. Images are then accessible through the FFIND web application, containing biological context-aware nomenclature, and are displayed by the interactive PathViewer client (Glencoe Software). Importantly, the native OMERO Plus web interface is bypassed and replaced with the FFIND front-end, designed to be fast and intuitive to use.

Developing Pancreatlas required interaction with software companies and exploration of open-source options to evaluate data management solutions that best aligned with our specific project objectives. While the popular and opensource OMERO system^6,7^ leverages Bio-Formats^8^ to access and integrate 150+ image formats and is sufficient for many basic image management needs, the imaging modalities we anticipated for Pancreatlas were already moving rapidly toward multiplexed systems like imaging mass cytometry (IMC)^9^ and co-detection by indexing (CODEX)^10^. Thus, we opted for OMERO Plus, which offers support and interactivity for images with 30-40 channels through PathViewer rather than OMERO’s standard image viewer (see **Table S1**).

PathViewer is particularly suited for appreciating images at both macro- (tissue architecture) and micro- (cellular) resolution, and the ability to hide and view individual channels or groups of channels also make it an ideal environment to share multiplexed imaging data that is laborious to store, transfer, and display in static environments. Moreover, the latest release (PathViewer 3) accommodates side-by-side viewing of multiple images, which we are currently working to integrate into our platform so that pre-curated sets of images can be loaded as a “single click” from our front-end site. This will be crucial to illustrating age- or disease-specific phenotypes; for example, users could have the option to view a particular image with a matched “control,” or to view the same marker combination at predefined age intervals to illustrate developmental changes. With an emphasis on adopting multiplexed imaging technologies, our group concluded that PathViewer was the most flexible, feature-rich browser available for image viewing and sharing in a meaningful biological context without requiring lengthy data downloads or software installation.

### Information Technology (IT) Infrastructure for FFIND and Pancreatlas

To run FFIND, the minimal infrastructure required is a modern client-based web browser and a single Linux-based server on which to install the FFIND API. It is up to developers to connect FFIND to appropriate data sources as well as integrate data viewers. For Pancreatlas, our infrastructure includes (1) a large virtualized server (CentOS7) to host the OMERO Plus server (eight-core Intel® Xeon® 2.1GHz, 32GB RAM); (2) several virtualized servers (each dual-core Intel® Xeon® 2.7GHz, 2GB RAM) for API and application development and hosting (RedHat Enterprise Linux 7); (3) on-site imaging repositories (Aperio eSlide Manager, Leica Biosystems; direct file high-performance storage for non-managed imaging data); (4) a cloud-hosted Laboratory Information Management System (LIMS) that contains tissue inventory and pancreas donor metadata; and (5) various web services used to connect and monitor overall architecture. The virtualization stack used provides scalable resources, including memory, processing capacity, and storage. Hence, we will be able to scale Pancreatlas without complicated and costly migrations due to hardware replacements. The bulk of our resources are co-located on the same 10Gb ethernet redundant network, and IT management is provided by several institutional support groups at Vanderbilt University and Vanderbilt University Medical Center.

### Testing and Monitoring

Testing and monitoring of our FFIND components and the Pancreatlas platform are achieved through automated code testing (Gitlab Continuous Integration), human-based functional testing (individual), focus group feedback sessions, local (Splunk) server monitoring, and remote service monitoring (Uptime Robot). We have load-tested 50 simultaneous users in our current hardware configuration to ensure reliable image viewing, and plan to continue testing performance under increased usage based on data collected through Google Analytics.

### Imaging Data and Metadata Imports into Pancreatlas

To manage the import and metadata annotations of our imaging data in Pancreatlas, we leveraged the out-of-the-box image import solution provided by OMERO Plus. We then customized this workflow to allow data curators/scientists – those individuals with domain knowledge and expertise in imaging data, including its acquisition and underlying biology – to work in a familiar spreadsheet environment (**Figure 2A**). Going by individual image or in sets of images, curators can annotate descriptive and structural metadata based on predefined, controlled vocabularies, and can select clinical donor attributes, which in our case were obtained from our cloud-based LIMS.

**Figure 2.**
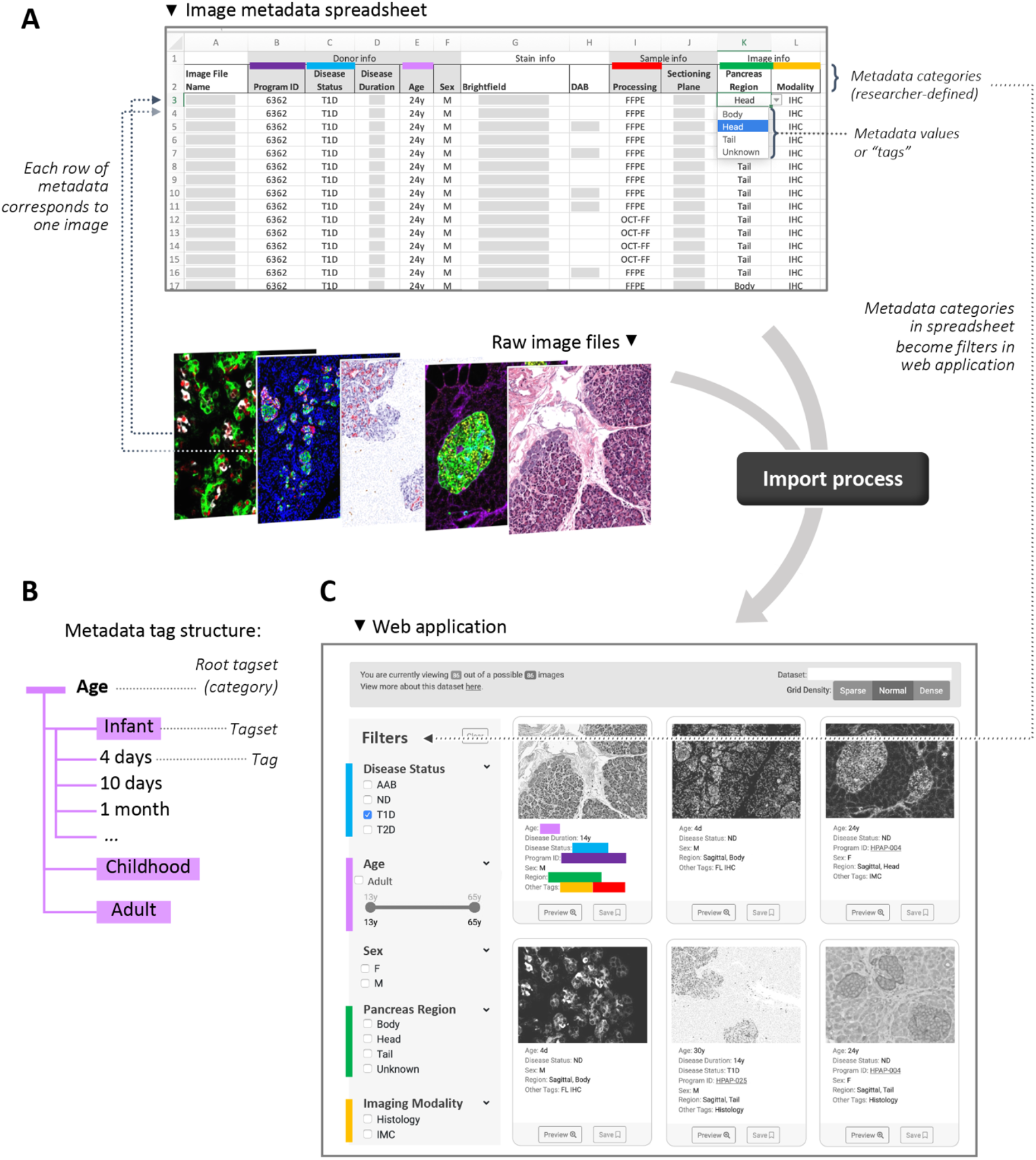
Pancreatlas metadata import process using FFIND infrastructure. **(A)** Data curators input various clinical and experimental details (metadata) for each data object, with metadata categories defined by the curator. This spreadsheet, a .csv file, directs automatic import of both raw data (shown here: images in Pancreatlas) and metadata. **(B)** During import, metadata are automatically parsed as “tags” assigned to each data object (tag = value from one cell of that object’s metadata row in A). If necessary, tags (shown here: 4 days, 10 days, 1 month) can be grouped into a broader tagset (shown here: Infant) to streamline FFIND’s filtering feature. **(C)** Data previews and metadata tags are retrieved through the custom web application. In the left column, simple filtering options are generated from metadata categories, represented by colored bars that correspond to those in panel A.

Once curators submit a data spreadsheet, the import mechanism within OMERO Plus validates the tabular data, ensures metadata completeness and image accessibility, and then proceeds to process images remotely and store newly created pyramidal image data within OMERO’s filesystem. This process concurrently populates several objects within OMERO’s PostgreSQL relational database^6^ and associates all of the imported annotations as object descriptors or key-value pairs. The FFIND filtering algorithm was configured to automatically parse predefined key-value pairs into tags or tagsets, thus enabling construction of a front-end filtering menu with biological relevance (**Figure 2, B-C**). After testing and necessary data refinements, an image collection is made public and displayed within the Pancreatlas front-end web application.

### Filtering Algorithm for Pancreatlas Image Data

One major design challenge of the FFIND platform was creating a system that allows for an arbitrary number of filters to be specified, based on the metadata tags and tagsets associated with FFIND data (**Figure 2B**). We define metadata tags as the specific value(s) of a metadata category that apply to a given piece of data (*e.g*., 5 years would be the metadata tag for the donor age category). A tagset refers to a collection of tags or tagsets that describe a category – for example, when considering donor age, age values can be grouped into specific stages of life (infant, childhood, adult, etc.), with each phase defined by the certain values/tags it includes (*e.g*., infant is 3 months to 24 months; childhood is 2 years to 10 years; adult is over 10 years). In this scenario, FFIND contains a root (broad) tagset of “age” which then contains three child tagsets of “infant”, “childhood”, and “adult,” each of which hosts a set of distinct age tags. In building our filtering algorithm, we aggregated all root tagsets into a single tree data structure to represent all possible filters for a given dataset. Using a depth-first traversal of this tree, we added logic for toggling filters on and off and updating the resulting list of matching images in the user interface. Importantly, this filtering component is connected directly to incoming URL query parameters, such that constructing complex filtering URLs is trivial and mirrors API calls.

### Pancreatlas Image Collections and Browsing Features

Pancreatlas utilizes FFIND’s web interface where data objects (images) are grouped into collections to seamlessly deliver contextual information, allowing users to apply filters and launch individual images in an interactive viewer (**Figure 3**). Importantly, these collections provide curated points of entry to an otherwise large quantity of images. In the default (grid) view of each collection (**Figure 4**), users can select and filter images by attributes of interest using various user interface (UI) widgets, with dynamic retrieval of applicable image previews. The user can then enlarge each image to reveal relevant metadata (donor age, gender, markers visualized, etc.) before they launch the full image viewer. In parallel, a matrix view allows selection of two attributes (*e.g*., age and gender) displayed in rows and columns, with each intersection populated by available images that meet the respective attribute values. These different viewing options, along with other key features of the web interface, are summarized in **Table 1**.

**Table 1.** Key features of FFIND and Pancreatlas.

| Feature | Description | Purpose/Significance | Figure(s) |
| --- | --- | --- | --- |
| *Data organization* |  |  |  |
| **Collections** | Datasets; the *Collections* page lists all available datasets, and each collection also has a dedicated page describing relevant background and details | Provides digestible overview of data breadth; gives project-specific context | 3A |
| **Multiple viewing modes** | <u>Grid view (default)</u>: data displayed as thumbnails with minimal descriptors (for easy browsing)<br><u>Temporal view</u>: data grouped by curated age ranges (for data that is temporal in nature)<br><u>Matrix view</u>: data grouped by user-selected attributes (e.g., age and sex), displayed in a manner that easily identifies whether data exists for specific attribute combinations (e.g., 6 years and male, 4 years and female, etc.) | Supports variable needs of user base; highlights key differences between data collections | 2C, 3B |
| **Data filtering** | Provides a flexible way to refine the data list into a subset of interest, allowing users to check boxes from lists of attribute options | Familiar interface (typical of eCommerce); intuitive to end users | 2C, 3B |
| *Data presentation* |  |  |  |
| **Thumbnails** | Image associated with each data object (in the case of Pancreatlas, an image) that highlights key features | Provides visual identifier for each data object | 3B, 4C |
| **Preview mode** | Enlarges thumbnail and lists additional metadata (tissue, experimental attributes) | Offers increased level of context while preserving filtered results | 4D |
| **Interactive browser (e.g., PathViewer)** | In the case of Pancreatlas, a web-based (HTML5) client by OMERO Plus that supports channel interactivity (toggle on/off, change colors, change range), accommodates 30+ channels within a single image, and offers multi-resolution zoom (pre-processed pyramidal data) | Allows users to view images in a web browser, from anywhere, without downloading any data; multi-resolution zoom enables appreciation of cellular scale within whole organ context | 3C-D |
| *General browsing* |  |  |  |
| **Bookmarking** | “Save” buttons on data cards/previews let the user add data to a bookmarked collection which persists over multiple sessions | Enables quick reference; users can build and share custom lists via unique URL | 4D |
| **Nomenclature page** | Lists all metadata terms, defining their biological relevance where applicable and providing diagrams to aid data interpretation | Encourages fieldwide adoption of metadata standards | 4E |

**Figure 3.**
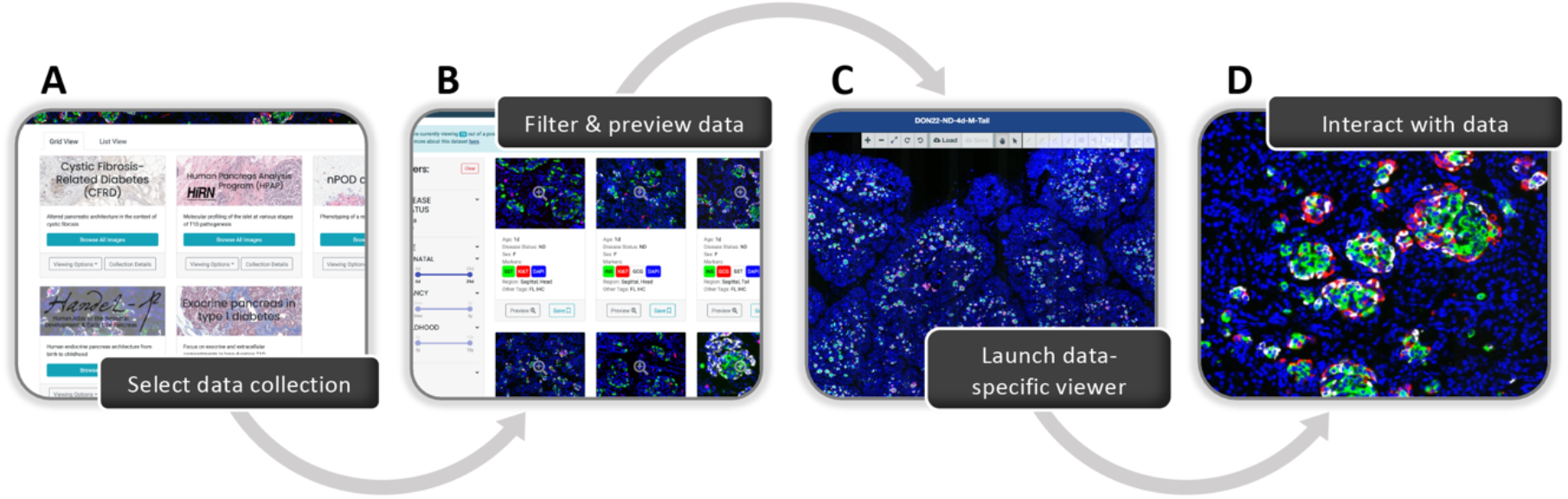
Stages of user-directed navigation via custom user interface in Pancreatlas. Shown are components of data navigation in Pancreatlas: **(A)** data collections page, **(B)** default grid view displaying data snapshots for a single collection, and **(C-D)** interactive web client for image browsing (PathViewer). For more detailed description of the user interface, see **Table 1**.

**Figure 4.**
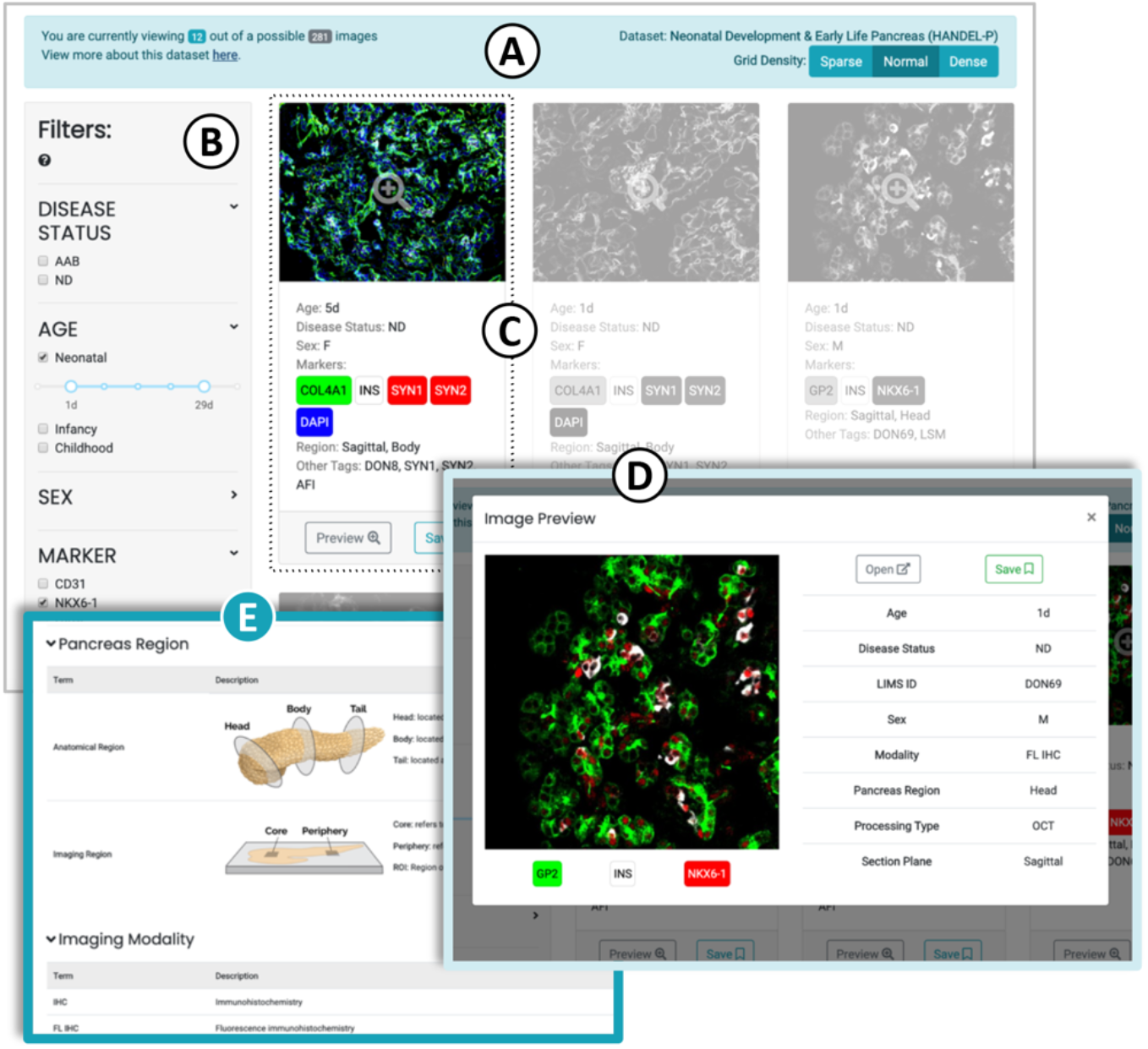
Image viewing within a selected Pancreatlas image collection. The default view features a data grid with navigation toolbar **(A)**, a filtering menu **(B)**, and small cards summarizing each data object (here, an image) **(C)**. Checking boxes in the filtering menu automatically repopulates cards only with data meeting the user-defined criteria. Clicking a data card launches a pop-up preview **(D)** that features an image thumbnail, experimental conditions, and other characteristics. From here, users can click on the thumbnail or *Open* button to launch PathViewer, or they can “save” the image to view later. Attribute terms in the filtering menu are defined on the Nomenclature page **(E)**, along with useful diagrams to aid the user in interpreting images.

To allow users to flag images of interest, the FFIND front-end has a bookmarking system where users can “save” images and display them in a session- and user-specific collection, as well as generate a shareable URL to return to the bookmarked images later. FFIND maintains this image list in a manner readable to the entire web application, enabling bookmarked images to be accessed in multiple parts of the application. The list is stored in the root application component of the React application and made accessible through callback functions in child components.

### Pancreatlas as an Image Publishing Platform

Five image collections have been made available in version 1.2 of Pancreatlas, each with a context-specific group of images and annotations (**Table 2**). These collections were compiled from various recent publications^11–15^ and pancreas phenotyping projects^16^ to illustrate significant pancreas-related biological processes or disease states. The imaging modalities range from conventional histology to multiplexed markers visualized by imaging mass cytometry.

**Table 2.** Image collections available in Pancreatlas.

| Name, description, & URL | # of images | Image type(s) | Reference |
| --- | --- | --- | --- |
| **Cystic Fibrosis-Related Diabetes (CFRD)**<br>Altered pancreatic architecture in the context of cystic fibrosis<br>http://www.pancreatlas.org/datasets/410/overview | **86 total**<br>(31 ND, 10 CF, 44 CFRD) | Whole-slide scans<br>(H&E, IHC, FL IHC) | Hart et al. 2018<sup>11</sup> |
| **Exocrine pancreas in type 1 diabetes - Focus on exocrine and extracellular compartments in long-duration T1D**<br>https://pancreatlas.org/datasets/703/overview | **135 total**<br>(73 ND, 62 T1D) | Whole-slide scans<br>(H&E, IHC, FL IHC) | Wright et al. 2020<sup>15</sup> |
| **Neonatal Development & Early Life Pancreas (HANDEL-P)**<br>Human endocrine pancreas architecture from birth to childhood<br>http://www.pancreatlas.org/datasets/531/overview | **281 total**<br>(274 ND, 7 AAB) | Whole-slide scans<br>(FL IHC), confocal<br>(FL IHC), CODEX | Manuscript in preparation |
| **Human Pancreas Analysis Program (HPAP)**<br>Molecular profiling of the islet at various stages of T1D pathogenesis<br>http://pancreatlas.org/datasets/508/overview | **303 total**<br>(129 ND, 77 T1D, 28 T2D, 69 AAB) | Whole-slide scans<br>(H&E), IMC | Wang et al. 2019<sup>9</sup>,<br>Kaestner et al. 2019<sup>16</sup> |
| **Network for Pancreatic Organ Donors with Diabetes (nPOD) case #6362**<br>Phenotyping of a recent-onset T1D donor<br>http://pancreatlas.org/datasets/525/overview | **39 total**<br>(all T1D) | Whole-slide scans<br>(H&E, IHC),<br>confocal (FL IHC) | Jackson et al. 2017<sup>12</sup>,<br>Canzano et al. 2018<sup>13</sup>,<br>Beery et al. 2019<sup>14</sup> |

AAB, autoantibody positive; CODEX, co-detection by indexing; CF, cystic fibrosis; FL, fluorescence; H&E, hematoxylin & eosin; IHC, immunohistochemistry; IMC, imaging mass cytometry; ND, non-diabetic; T1D, type 1 diabetes; T2D, type 2 diabetes.

One limitation of traditional scientific publications, even those available online, is the constrained space to present primary data. For example, in a recent study of cystic fibrosis-related diabetes (CFRD) by Hart and colleagues^11^, 12 images were published in the main paper with another 7 in the supplement – however, the reported analyses utilized more than 80 images. This full image set has been made available via Pancreatlas as a disease-specific collection (*CFRD*, **Table 2**), and it offers the added advantage for users to navigate around large tissue areas, zoom in on regions of interest, and interact with the data in a way that is not possible with traditional publication formats. Another project that highlights the value of flexible spatial resolution (*i.e*., viewing cells at high magnification but also retaining the large-scale tissue context) is the investigation of processes governing human pancreatic development. In an effort to gain insight into possible triggers of type 1 diabetes, a forthcoming study from our group closely examines islet composition and architecture from birth to ten years of age. Pancreatlas currently houses 281 images from this study (*HANDEL-P*, **Table 2**), many of which are whole-slide scans measuring up to 900 megapixels, or 30,000 pixels in both dimensions. Access to such high-resolution data is critical to appreciating the spatiotemporal context of pancreas and islet development; images provide detailed information of small islet structures (100-200 μm diameter) within the landscape of entire pancreatic cross-section (10-20 cm^2^). As highlighted in **Table 1**, the dynamic interactivity facilitated by PathViewer is critical to understanding development on a whole-organ scale.

## Discussion

The development of FFIND and Pancreatlas was stimulated by our group’s desire to effectively share imaging data from human tissue along with the associated phenotypic or clinical traits, where applicable. Image databases are notoriously challenging and laborious to construct due to the large file sizes and the need to assimilate multiple imaging modalities and data formats. Whereas innumerable solutions have been developed for sequencing datasets (*Nucleic Acids Research* publishes an annual Database Issue and maintains an online list of thousands of molecular biology databases^17^), platforms for imaging data lag noticeably behind. The structural imaging community has addressed this shortage by building repositories with support from the European Molecular Biology Laboratory-European Bioinformatics Institute (EMBL-EBI); the Electron Microscopy Data Bank (EMDB)^18^ and Electron Microscopy Public Image Archive (EMPIAR)^19^ provide 3D reconstructions and raw 2D data, respectively, for protein structures obtained through cryo-electron microscopy. Excitingly, the first public “added-value” bioimage databases have recently emerged: Image Data Resource^20,21^ (IDR) and the Systems Science of Biological Dynamics Database^22^ (SSBD). They accept submissions for reference datasets and provide substantial annotation and linkage to external resources. Pancreatlas joins these public resources as a bioimage database publishing reference datasets related to the biology and pathology of the pancreas. In the longer term, a common repository for all bioimage datasets related to published studies will be required – development of the BioImage Archive (EBI) is a first step towards this goal^23^.

Like other added-value databases, Pancreatlas aims to provide key datasets that can be referenced and/or reanalyzed thanks to their detailed metadata annotations. However, our platform also offers a visually- and organizationally-cohesive environment in which to explore disparate data collections. Pancreatlas represents a departure from many existing organ-specific resources that launch multiple different tabs, viewers, and/or annotation schemes depending on the data being explored. While different data types will always require specialized clients or browsers, the FFIND interface equips users with an understanding of all available data from a common “jumping-off” point. IDR maintains a consistent browsing experience through modular collections similar to those in Pancreatlas; however, IDR users interact directly with the standard OMERO web client, whereas Pancreatlas users instead navigate an intuitive FFIND web interface, which masks the underlying OMERO Plus server and connects the user to a rich image viewing client, PathViewer.

As we highlight in **Figure 1**, FFIND and its implementation, Pancreatlas, hold great potential to enrich existing databases or repositories by adding flexible, user-friendly interfaces to broaden audience and impact. We feel that FFIND represents a worthy model for data scientists and developers in this capacity, as it is a solution that can be re-used for any domain. Available as open-source software, FFIND can be downloaded and installed in less than five minutes, and it provides a simple approach to tie into existing APIs and data viewers without requiring refactoring of large or complex codebase. Unlike other “atlas” or “mapping” groups and consortia, we have only a small team that handles data curation, annotation, infrastructure, and development – yet despite a smaller size and funding level, Pancreatlas is still positioned to benefit human disease research and eventually human health. We believe that other small groups needing a solid foundation to begin or augment a data sharing platform will find FFIND to be an accessible, cost-sensitive solution with exceptional opportunity to cater to domain-specific needs.

As an example of FFIND’s data-agnostic use, we are also investigating how to leverage transcriptome analysis results available through the RNASeq-er API, developed by EMBL-EBI^24^. This web service provides access to the results of standardized alignments as well as gene and exon expression quantification of all public bulk (and eventually single cell) RNA sequencing experiments stored at the European Nucleotide Archive. A curated set of studies could be assembled and retrieved by FFIND, having been configured to incorporate associated ontology-based terms and tagsets into its filtering system, and then connected to a suitable viewer, such as DEBrowser (an R Shiny app). DEBrowser would be used for pre-analyzed comparisons or to perform new differential gene expression analysis in a web browser, using well-established methods^25^. To the best of our knowledge, this framework could be configured in a short period of time with minimal development – allowing instead for focus on curation and exploratory data analysis.

In the case of Pancreatlas, our efforts thus far have focused on providing an intuitive, centralized platform to facilitate sharing of high-value imaging data sets. By bringing data from different programs and initiatives under one “roof” and investing time in integrating with other databases, we hope to accelerate and communicate molecular discoveries related to human pancreas biology. We envision that Pancreatlas image collections will inform both study design (*e.g*., providing reference data for various stages of pancreatic development), as well as connect research fields (*e.g*., diabetes and pancreatic cancer) to identify collaborations and interdisciplinary pursuits that integrate cellular-resolution data within the context of whole-organ architecture. We are also committed to facilitating data exploration from high-impact publications in pancreas research, thus reducing the burden on researchers (or journals) to navigate general repositories that necessarily lack field-specific annotations. In fact, FFIND offers a potential solution to any researcher who has data in a public repository, providing them a framework for a customizable, public-facing web resource with minimal investment. As shown in Pancreatlas, FFIND can easily associate datasets with a publication DOI, and we hope ongoing conversations with leadership in the scientific publishing community will result in partnerships to broaden manuscript impact and promote cross-disciplinary discoveries. Hopefully, this will empower the creation of more added-value resources referencing data from centrally maintained repositories.

### Future Work

We are continuing to improve the FFIND technology platform that supports Pancreatlas, including IT infrastructure improvements (Amazon Web Services deployment), back-end improvements (abstraction, documentation), and front-end improvements (user interface consistency, device-agnostic usability, performance improvements, and error detection). Over the next year we will be adding several new features, including a global browsing tool that will give users the option to access data across all collections simultaneously. We will also extend the FFIND API to provide a public and documented API, in order to allow other groups to collage datasets and then request those for download. Simultaneously, we hope to establish partnerships with existing image data repositories such as IDR for long term data deposition and sustainability, and from which interested parties could easily download our data. For Pancreatlas specifically, we will be updating our nomenclature to reference ontology-based terms (Experimental Factor Ontology^26^, Human Disease Ontology^27^, Ontology for Biomedical Investigations^28^, Measurement Method Ontology^29^, Foundational Model of Anatomy Ontology^30^, and Biological Imaging Methods Ontology). We plan to leverage new PathViewer features like multi-image views in order to create more focused stories that will provide curated knowledge to selected data. We will also continue to promote inter-program connectivity, as we have done with other pancreas-specific resources like the Human Pancreas Analysis Program (HPAP)^16^ and the Integrated Islet Distribution Program (IIDP)^31^. Finally, we are cognizant of the need to communicate biomedical research beyond the academic community, and we are exploring ways to make Pancreatlas more approachable to those without domain-specific knowledge. For example, the recently developed tool Minerva from Rashid and colleagues^32^ represents an exciting opportunity to incorporate narrative guides into the presentation of histological images. Software tools such as this one could be utilized to highlight unique aspects of individual datasets and image collections.

### Sustainability

Longevity is a universal challenge for database platforms; even successful, established resources require sustained funding to maintain and/or expand infrastructure, curate data, and provide users with tools necessary for the ever-evolving analysis paradigms to understand large datasets^33–35^. With Pancreatlas, we already track usage by collecting voluntary information from users (*e.g*., type of institution with which the user is associated) and we are collaborating with others in the community to leverage national federal funding as well as private support. One of FFIND’s advantages is its ability to be deployed and maintained by a small team, relying on local, open-source, and/or commercial back-ends for data storage. Should Pancreatlas need to shut down at some point, all of its metadata (stored in a portable and open-source PostgreSQL database) and images (supported by Bio-Formats) could easily be deposited to open-access repositories like Zenodo, Dryad, or others.

Ideally, we envision Pancreatlas data being housed in public repositories such as IDR^20^. While running our web interface off of IDR (an EBI-based resource) would pose logistical and performance challenges, there are no technical hurdles to storing Pancreatlas images in a data- or topic-agnostic repository or database. We welcome the opportunity to collaborate with other database teams and are committed to working and promoting the application of FAIR principles^36^ to bioimaging data. Building added-value public resources for scientists requires continued conversation and collaboration among researchers, scientific journals, and biomedical research funders^21,37^. We view both FFIND and Pancreatlas as living platforms that will evolve with continued collaboration and input from the broader community, and we look forward to partnering with other scientists to advocate for the value of data sharing through accessible and thoughtfully designed database solutions.

#### “Bigger Picture”

Scientists need cost-effective yet fully-featured database solutions that facilitate large dataset sharing in a structured and easily digestible manner. Flexible Framework for Integrating and Navigating Data (FFIND) is a data-agnostic, scalable web interface that is designed to integrate with existing databases and data-browsing clients while offering an intuitive front-end website. We used FFIND to build Pancreatlas, an online imaging resource containing datasets linking imaging data with clinical data to facilitate advances in understanding of diabetes, pancreatitis, and pancreatic cancer. FFIND architecture, which is available as open-source software, can be easily adapted to meet other field- or project-specific needs; we hope it will help data scientists reach a broader audience by reducing the development lifecycle and providing familiar interactivity in communicating data and underlying stories.

## Experimental Procedures

### Source Code Availability

FFIND is available as open-source software; please visit https://github.com/Powers-Brissova-Research-Group/FFIND for repository access and https://powers-brissova-research-group.github.io/FFIND/ for a deployed instance. FFIND and Pancreatlas are committed to sharing data according to FAIR principles.

## Acknowledgements

We are extremely grateful to Jason Swedlow, Chris MacLeod, and Emil Rozbicki of Glencoe Software, who provided crucial technical and troubleshooting support and enthusiasm for our project. We also thank Dr. Swedlow for his critical reading of this manuscript. This work was performed using resources and/or funding provided by The Leona M. and Harry B. Helmsley Charitable Trust, by the NIDDK-supported Human Islet Research Network (RRID:SCR_014393; http://hirnetwork.org; UC4 DK104211, DK108120, DK112232, DK123716), by DK106755 and DK20593, and by the Department of Veterans Affairs (BX000666).

## Author Contributions

Conceptualization, all authors; Methodology, D.C.S., J.M., M.B., and J.P.C.; Software, J.M. and J.P.C.; Resources, I.K., M.L.B., M.Y., and M.A.; Data Curation, D.C.S. and M.L.B.; Writing – Original Draft, D.C.S., J.M., M.B., and J.P.C.; Writing – Review & Editing, all authors; Supervision, M.A., A.C.P., M.B., and J.P.C.; Project Administration, I.K., M.L.B., and M.Y.; Funding Acquisition, M.A. and A.C.P.

## Supplemental Information

**Table S1.** Comparison of OMERO/OMERO.iviewer and OMERO Plus/PathViewer for image viewing.

| Feature or Component | OMERO.iviewer (OMERO) | PathViewer (OMERO Plus) |
| --- | --- | --- |
| Multi-image views | Supported in newer versions | Supported and tailored to digital pathology workflows |
| Loading images | Support for multi-resolution, tiled images | Support for multi-resolution, tiled images<br>• Support for acceleration through tile microservice<br>• Support for selecting tile sizes and compression parameters<br>• Support for image flipping through tile microservice<br>• Predictive preloading of images |
| Annotation tools | Basic tools required for marking and annotating regions of interest (ROIs) | Extended features for ROI annotation, including improved freehand drawing and detailed rich formatted descriptions<br>• Rich formatted descriptions allowed at Image and Channel levels |
| Multichannel image support | Supported | Supported<br>• Ability to organize channels into custom-named groups and toggle on/off<br>• Intuitive design for adjusting brightness and contrast |
| Portability | Parametric URL available from right-click; status of ROI display not included | Parametric URLs to share exact viewing settings available from Location bar |
| Metadata | Basic metadata visible in separate Info tab; map annotations not supported | Annotations and map annotations viewable in Properties panel, along with image info and description |

Sources: Glencoe Software, Inc. (n.d.). *PathViewer*. http://www.glencoesoftware.com/products/pathviewer/features/ and University of Dundee & Open Microscopy Environment. (n.d.). *OMERO.iviewer*. http://www.openmicroscopy.org/omero/iviewer/

**Table S2.**
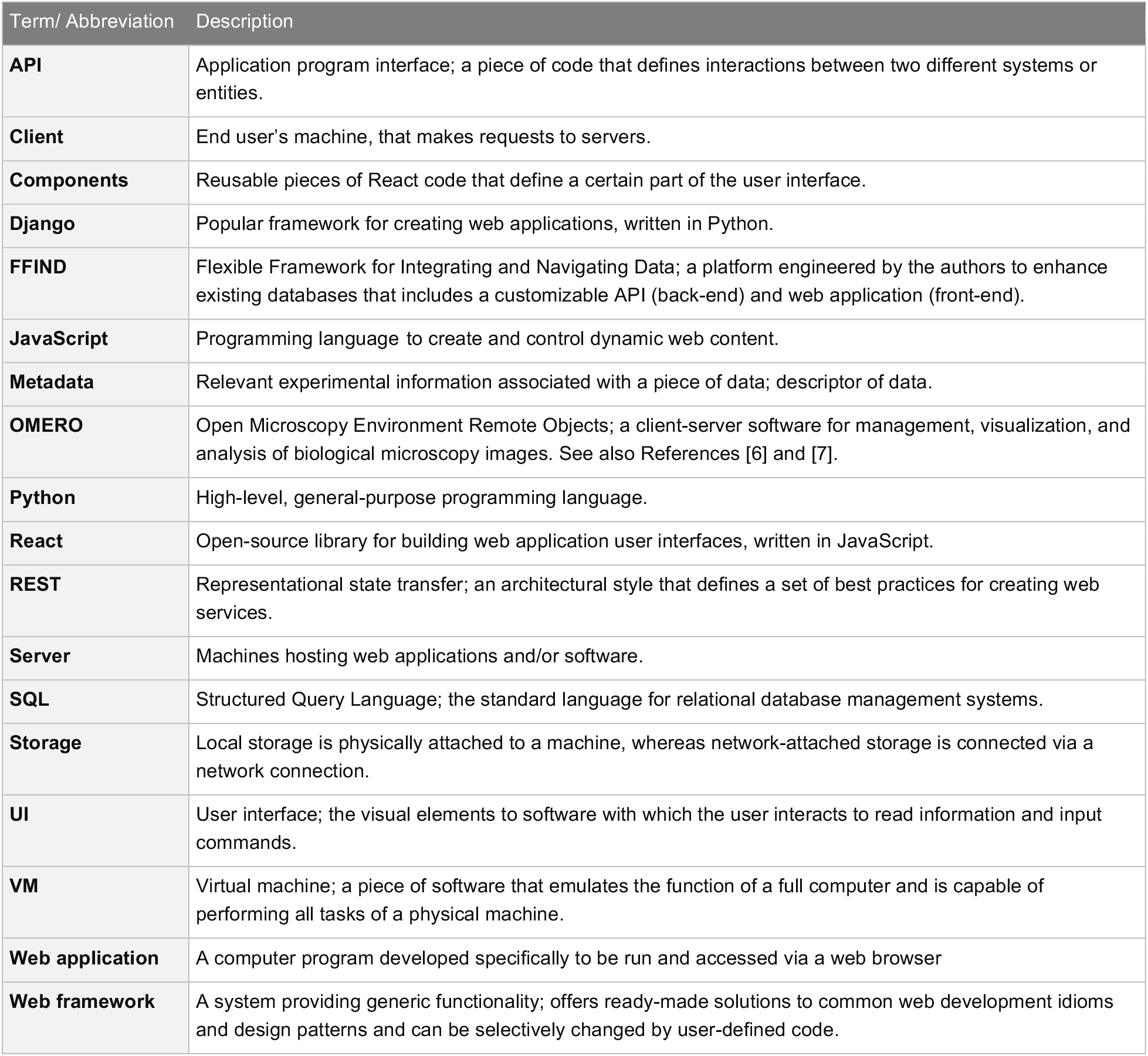
Glossary of abbreviations and technical terms.

| Term/ Abbreviation | Description |
| --- | --- |
| **API** | Application program interface; a piece of code that defines interactions between two different systems or entities. |
| **Client** | End user's machine, that makes requests to servers. |
| **Components** | Reusable pieces of React code that define a certain part of the user interface. |
| **Django** | Popular framework for creating web applications, written in Python. |
| **FFIND** | Flexible Framework for Integrating and Navigating Data; a platform engineered by the authors to enhance existing databases that includes a customizable API (back-end) and web application (front-end). |
| **JavaScript** | Programming language to create and control dynamic web content. |
| **Metadata** | Relevant experimental information associated with a piece of data; descriptor of data. |
| **OMERO** | Open Microscopy Environment Remote Objects; a client-server software for management, visualization, and analysis of biological microscopy images. See also References [6] and [7]. |
| **Python** | High-level, general-purpose programming language. |
| **React** | Open-source library for building web application user interfaces, written in JavaScript. |
| **REST** | Representational state transfer; an architectural style that defines a set of best practices for creating web services. |
| **Server** | Machines hosting web applications and/or software. |
| **SQL** | Structured Query Language; the standard language for relational database management systems. |
| **Storage** | Local storage is physically attached to a machine, whereas network-attached storage is connected via a network connection. |
| **UI** | User interface; the visual elements to software with which the user interacts to read information and input commands. |
| **VM** | Virtual machine; a piece of software that emulates the function of a full computer and is capable of performing all tasks of a physical machine. |
| **Web application** | A computer program developed specifically to be run and accessed via a web browser |
| **Web framework** | A system providing generic functionality; offers ready-made solutions to common web development idioms and design patterns and can be selectively changed by user-defined code. |

